# Structure of human phagocyte NADPH oxidase in the resting state

**DOI:** 10.1101/2022.10.04.510768

**Authors:** Rui Liu, Kangcheng Song, Jing-Xiang Wu, Xiao-Peng Geng, Liming Zheng, Xiaoyin Gao, Hailin Peng, Lei Chen

**Author notes:** These authors contributed equally to this work (Rui Liu, Kangcheng Song). Corresponding author: Lei Chen.

## Abstract

Phagocyte oxidase plays an essential role in the first line of host defense against pathogens. It oxidizes intracellular NADPH to reduce extracellular oxygen to produce superoxide anions for pathogen killing. The resting phagocyte oxidase is a heterodimeric complex formed by two transmembrane proteins NOX2 and p22. Despite the functional importance of this complex, its structure remains elusive. Here we reported the cryo-EM structure of the human NOX2-p22 complex in nanodisc in the resting state. The structure shows that p22 is formed by four transmembrane helices and interacts with NOX2 through its M1 and M4 helices. Hydrophobic residues on M3, M4, and M5 of NOX2 contribute to the complex formation. Structural analysis suggests that the cytosolic factors activate the NOX2-p22 complex by stabilizing the dehydrogenase domain (DH) in a productive docked conformation which is efficient for electron transfer between DH and the transmembrane domain.

## Introduction

NADPH oxidases (NOX) are membrane-bound redox enzymes ^1^. They transfer electrons from intracellular NADPH to extracellular oxygen to generate reactive oxygen species (ROS), including superoxide anions and hydrogen peroxide ^1^. There are seven NOX members identified in humans, including NOX1-NOX5 and DUOX1-2 ^2^. They are involved in broad physiological processes, such as host defense, cell signaling, protein modification, and hormone synthesis ^1^.The mutations of genes encoding NOX proteins can cause human diseases such as immunodeficiency and hypothyroidism ^2^.

NOX2 is the key catalytic subunit of phagocyte oxidase, which is highly expressed in phagocytes and is essential for innate immunity ^3^. Resting phagocyte oxidase is a heterodimer composed of two multi-pass transmembrane proteins NOX2 (gp91^phox^) and p22^phox^ (p22) ^3^. Once invading pathogens are trapped inside the special membrane compartment named phagosomes, a phosphorylation-dependent signaling cascade activates the cytosolic factors of NOX2, including p47^phox^, p67^phox^, p40 ^phox^, and Rac ^3^. Activated cytosolic components dock onto and activate the NOX2-p22 heterodimer, which transfers electrons from intracellular NADPH to oxygen, causing the respiratory burst of phagocytes. The generated superoxide anions could kill the invading pathogens inside phagosomes ^3^. In accordance with the key physiological function of NOX2, the loss-of-function mutations of the *NOX2* gene lead to failures of clearance of invading pathogens due to low level of superoxide anions produced, causing chronic granulomatous disease (CGD)^4^.

Previous studies show that the catalytic core of NOX has a transmembrane domain (TMD) and a cytosolic dehydrogenase domain (DH)^5-7^. TMD has six transmembrane helices which coordinate two haems for electron transfer across the membrane ^5-7^. DH shares sequence homology to the ferredoxin-NADP reductase (FNR)^5-7^. DH domain is formed by the FAD-binding sub-domain (FBD) and NADPH binding sub-domain (NBD)^5-7^. The available crystal structure of the transmembrane domain (TMD) of homologous csNOX5 revealed the electron transfer pathway across the membrane and the oxygen substrate binding site ^5^. Recent cryo-EM structures of the DUOX1-DUOXA1 complex showed how the FAD cofactor and NADPH substrate are bound at the interface between the DH domain and TMD of DUOX ^6,7^. Moreover, the structure of human DUOX1-DUOXA1 in the high-calcium state revealed the conformation of the activated NOX^7^. These studies have provided instrumental information about the structure of NOX2. However, different from NOX5 or DUOX, the function of NOX2 protein absolutely requires an essential auxiliary subunit p22 ^2^. p22 harbors the docking site for the cytosolic component p47^phox^ protein which is crucial for the assembly of NOX2 cytosolic components. The loss-of-function mutations of p22 also lead to CGD ^4^, emphasizing its physiological importance. Notably, p22 can also form a complex with NOX1, NOX3, and NOX4 and is essential for the functions of these proteins. Prior to our current study, the structure of p22 is largely unknown due to the lack of homology models, and the topology of p22 is controversial because different studies suggest that p22 has 2 or 3, or 4 transmembrane helices ^4,8,9^. More importantly, how p22 interacts with NOX1-4 protein remains mysterious. Here we present the structure of the human NOX2-p22 complex in the resting state which uncovers the architecture of this important enzyme.

## Results

### Structure determination

To measure the superoxide anions released from the NOX2 enzyme, we converted the superoxide anion into hydrogen peroxide using SOD and used the Amplex Red assay ^7,10^ to measure the concentration of hydrogen peroxide produced in real-time (Figure 1A). To activate NOX2 in a cell-free system, we exploited a highly engineered chimeric protein named Trimera, which contains essential fragments of p47 ^phox^, p67 ^phox^, and Rac1 for NOX2 activation ^11^. Using this assay system, we found that co-expression of human NOX2 and C-terminally GFP-tagged human p22 could yield cell membranes that showed Trimera-activated superoxide anion production which is sensitive to DPI inhibition (Figure supplement 1A), suggesting a correctly assembled NOX2-p22 hetero-complex. However, subsequent purification and cryo-EM analysis of this complex only resulted in a medium-resolution reconstruction that was not sufficient for confident model building. We reasoned that the small size of the complex and the endogenous conformational heterogeneity of NOX2 DH might represent major obstacles to our structural studies. Therefore, we exploited the possibility of using a commercially available mouse anti-human NOX2 monoclonal antibody 7D5 ^12^ as a fiducial marker for cryo-EM studies. We produced the Fab fragment of 7D5 by papain digestion and bound the Fab with strep-tagged anti-mouse kappa chain nanobody TP1170 ^13^. We reconstituted the human NOX2-p22 hetero-complex in nanodisc and purified the NOX2-p22-7D5-TP1170 complex (Figure supplement 1B-C). The resulting quaternary complex showed robust Trimera-dependent DPI-sensitive superoxide production (Figure 1B-C), indicating it is a functional NOX2-p22 complex that is suitable for structural studies. Cryo-EM data collection of the protein sample absorbed on both graphene-oxide coated grids ^14^ and graphene-coated grids ^15^ allowed the structural determination of this asymmetric complex to an overall resolution of 3.3 Å (Figure supplement 2). However, the local map quality of the intracellular DH domain of NOX2 and the constant region of Fab (CH1) and TP1170 on the extracellular side was poor in the consensus map. Subsequent local refinement focusing on these two regions using mask1 and mask2 further improved their local map quality (Figure supplement 2). Therefore we merged the focused refined map into the consensus map to generate a composite map for model building and interpretation (Supplementary Table 1).

**Fig. 1.**
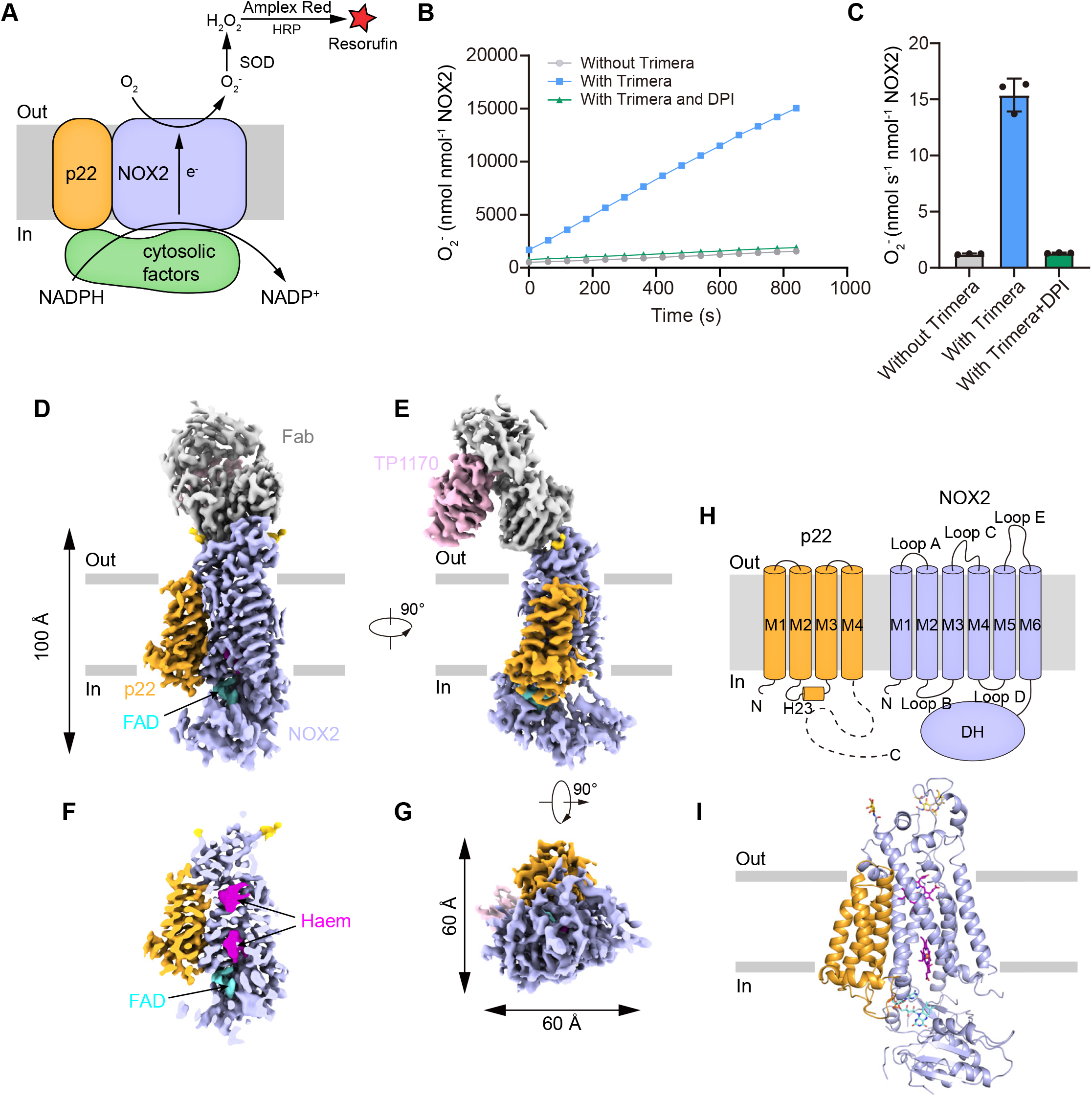
The structure of the human NOX2-p22 complex at the resting state A, Schematic of the NOX2 enzymatic assay. O_2_^-^ produced by NOX2 are converted into H_2_O_2_ by SOD. In the presence of H_2_O_2_, horseradish peroxidase (HRP) converts the nonfluorescent Amplex Red into resorufin, the fluorescence of which is measurable and proportional to the concentration of H_2_O_2_. B, The amounts of O_2_^-^ produced by NOX2-p22-7D5-TP1170 complex in nanodisc in the presence of Trimera are plotted versus time. The enzymatic system contains 0.36 nM NOX2, 33nM Trimera, and 10 μg/ml DPI, as indicated. The represented data are shown. DPI, diphenyleneiodonium, NADPH oxidase inhibitor. C, The rates of O_2_^-^ production in B are summarized. Data are shown as means ± standard deviations; N = 3 technical replicates. D, Side view of the cryo-EM map of NOX2-p22-7D5-TP1170 complex. The approximate boundaries of the phospholipid bilayer are indicated as gray thick lines. p22 and NOX2 are colored in orange and light blue. 7D5 and TP1170 are colored in grey and pink, respectively. E, A 90°-rotated side view compared to D. F, The cut-open view of the cryo-EM map of NOX2-p22. Haem and FAD is colored in magenta and cyan, respectively. G. A 90°-rotated bottom view of E. H, Topology of p22 and NOX2 subunits. Transmembrane helices are shown as cylinders, unmodeled disordered regions are shown as dashed lines. The phospholipid bilayer is shown as gray layers. DH, dehydrogenase domain of NOX2. I, Structure of NOX2-p22 complex in cartoon representation. The colors are the same as in D.

**Table 1.** Cryo-EM data collection, refinement, and validation statistics. The parameters for the Cryo-EM data collection, processing, and validation of the NOX2-p22-7D5-TP1170 complex are listed in the table.

| PDB ID<br>EMDB ID | NOX2-p22-7D5-TP1170<br>8GZ3<br>EMD-34389 |
| --- | --- |
| **Data collection and processing** |  |
| Magnification | 165,000 × |
| Voltage (kV) | 300 |
| Electron exposure (e<sup>-</sup>/Å<sup>2</sup>) | 37.6 |
| Defocus range (μm) | -1.5 to -2.0 |
| Pixel size (Å) | 0.821 |
| Symmetry imposed | *C1* |
| Initial particle images (no.) | 541,609 |
| Final particle images (no.) | 141,189 |
| Map resolution (Å) | 3.3<sup>a</sup>/3.8<sup>b</sup>/3.7<sup>c</sup> |
| FSC threshold | 0.143 |
| Map resolution range (Å) | 250-3.3 |
| **Refinement** |  |
| Initial model used (PDB code) | Alpha Fold 2 |
| Model resolution (Å) | 3.3 |
| FSC threshold | 0.143 |
| Model resolution range (Å) | 250-3.3 |
| Map sharpening *B* factor (Å<sup>2</sup>) | -156.3<sup>a</sup>/-213.2<sup>b</sup>/-193.9<sup>c</sup> |
| Model composition |  |
| Non-hydrogen atoms | 9,044 |
| Protein residues | 1,150 |
| Ligands | 6 |
| *B* factors (Å<sup>2</sup>) |  |
| Protein | 74.24 |
| Ligand | 58.47 |
| R.m.s. deviations |  |
| Bond lengths (Å) | 0.005 |
| Bond angles (°) | 0.685 |
| Validation |  |
| MolProbity score | 1.66 |
| Clashscore | 6.81 |
| Poor rotamers (%) | 0.00 |
| Ramachandran plot |  |
| Favored (%) | 95.89 |
| Allowed (%) | 4.11 |
| Disallowed (%) | 0.00 |

a, The values of consensus refinement.
b, The values of focused refinement of mask1 (CH1+TP1170).
c, The values of focused refinement of mask2 (DH).

The final structural model contains most residues of NOX2 and 3-136 of p22 (Figure 1, Figure supplement 4 and 5). The long cytosolic tail of p22 is invisible in the cryo-EM map due to its flexibility. The structure shows that p22 has a four-helix bundle architecture and binds to the NOX2 TMD from the side, forming a hetero-complex occupying 100 Å × 60 Å × 60 Å in the three-dimensional space (Figure 1 D-G). The NOX2-p22 complex structure allowed us to map the spatial positions of NOX2 and p22 mutations found in CGD patients (Figure supplement 6). 7D5 Fab recognizes a structural epitope on the extracellular side of NOX2 (Figure 1D-E).TP1170 binds to the kappa light chain constant domain of 7D5 Fab (Figure 1E), which is consistent with previous functional data ^13^. NOX2 requires cytosolic factors for activation ^3^. Because there are no cytosolic factors supplemented in the cryo-EM sample, the current structure represents the human NOX2-p22 complex in the inactive resting state.

### Structure of human NOX2

NOX2 resembles the canonical structure of NOX, which is consist of an N-terminal TMD and a C-terminal cytosolic DH domain (Figure 2A). The extracellular region is formed by Loop A, Loop C, and Loop E (Figure 2B, Figure supplement 4). Loop E sits on the top and there is an intra-loop disulfide bond (C244-C257) which stabilizes Loop E in a compact structure (Figure 2B). Notably, C244R^16^, C244S^17^, and C244Y^18^ mutations on NOX2 were previously identified in CGD patients (Figure supplement 6). These mutations likely affect the conformational stability of Loop E and thus the maturation of NOX2. We observed two N-link glycosylation decorations at N132 and N149 on Loop C (Figure 2B). Loop C and Loop E form the structural epitope that is recognized by 7D5 Fab (Figure 1D-E, Figure 2B). Loop A buttresses below Loop C and Loop E to support their conformation (Figure 2B). These three extracellular loops form a dome that caps above the outer haem bound in the TMD of NOX2 (Figure 2A-C). Highly conserved R54, H119, and the outer haem surround the oxygen-reducing center (Figure 2D). We found a hydrophilic tunnel that connects the extracellular environment and the oxygen-reducing center (Figure 2C).

**Fig. 2.**
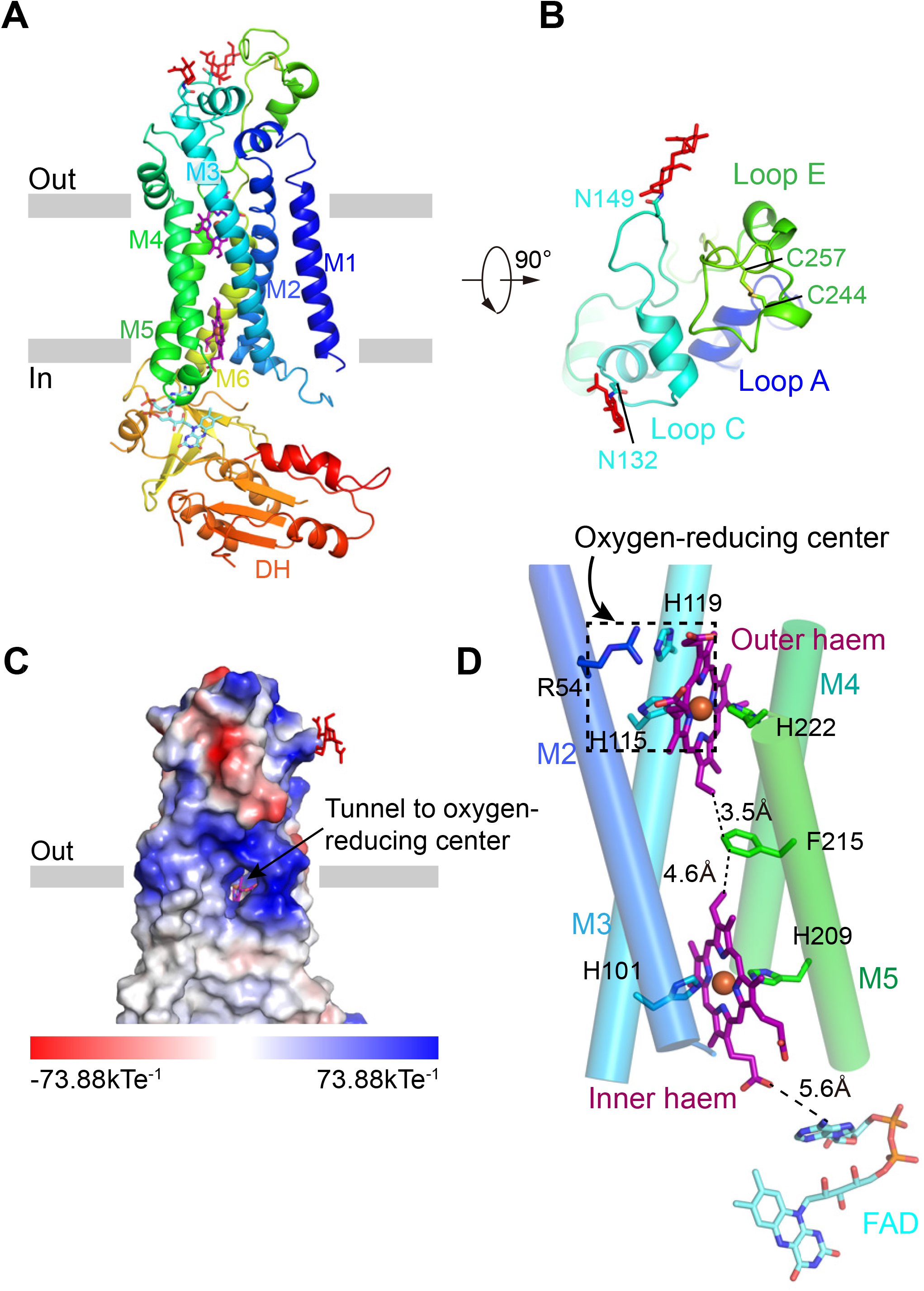
The structure of human NOX2 A, Structure of NOX2 in cartoon representation, and is colored in the rainbow pattern (N-terminus is blue and C-terminus is red). The phospholipid bilayer is shown as gray layers. B, A 90°-rotated top view of A. The disulfide bonds and glycosylation are shown as sticks. Three extracellular loops are indicated. C, Surface electrostatic potential distribution of NOX2. The tunnel to the oxygen reducing center is indicated with an arrow. Haem and glycosylation are shown as sticks. The phospholipid bilayer is shown as gray layers. D, The components on the electron transfer pathway of NOX. The edge-to-edge distances between adjacent components are shown in dashes. The oxygen reducing center is indicated as a dashed rectangle. The ligands are shown as sticks. The ferric ions are shown as spheres. The helices are shown as cylinders. The colors are the same as in A.

This tunnel might represent the putative pathway for the entry of oxygen substrate and release of superoxide anions. The outer haem, inner haem, and F215 between them form the electron-transferring pathway across the membrane (Figure 2D). Outer haem is coordinated by H115 on M3 and H222 on M5. Inner haem is coordinated by H101 on M3 and H209 on M5 (Figure 2D). Below the TMD, we observed a strong density of FAD which binds in the FBD of DH (Figure 2D and Figure supplement 3D). But the density of NBD is poor even after focused refinement, likely due to its high mobility (Figure supplement 2E,F and Figure supplement 3). We did not observe the density of NADPH (Figure supplement 3), although we have supplemented NADPH into the cryo-EM samples. The structure of NOX2 allowed us to measure the edge-to-edge distances as 3.5 Å between the outer haem and F215; 4.6 Å between F215 and the inner haem; 5.6 Å between the inner haem and FAD (Figure 2D).

### Structure of human p22

The structure of p22 is consist of four tightly packed transmembrane helices (Figure 3). An amphipathic helix H23 on the M2-M3 linker floats on the inner leaflet of the membrane (Figure 3A). Different from common TMD structures which are usually formed by hydrophobic residues, a polar interaction network, formed by side chains of N11 on M1, E53 on M2, R90 and H94 on M3, and Y121 on M4 is caged inside the TMD of p22 (Figure 3A). These residues are highly conserved in p22 (Figure supplement 5). Moreover, they are CGD mutation hotspots. R90Q ^19^, R90W^20^, E53V^21^, and H94R^19^ mutations of p22 have been identified in CGD patients (Figure supplement 6), likely because the polar interaction network formed by these residues is important for the stability of p22. Below the TMD, the 128-136 segment of p22 extends out of M4 and binds into a crevice shaped by H23 and M2-H23 loop on the M2-M3 linker (Figure 3B). Such interactions might position the p22 tail in a suitable pose that is ready for p47^phox^ recruitment and subsequent assembly of other cytosolic factors during NOX2 activation.

**Fig. 3.**
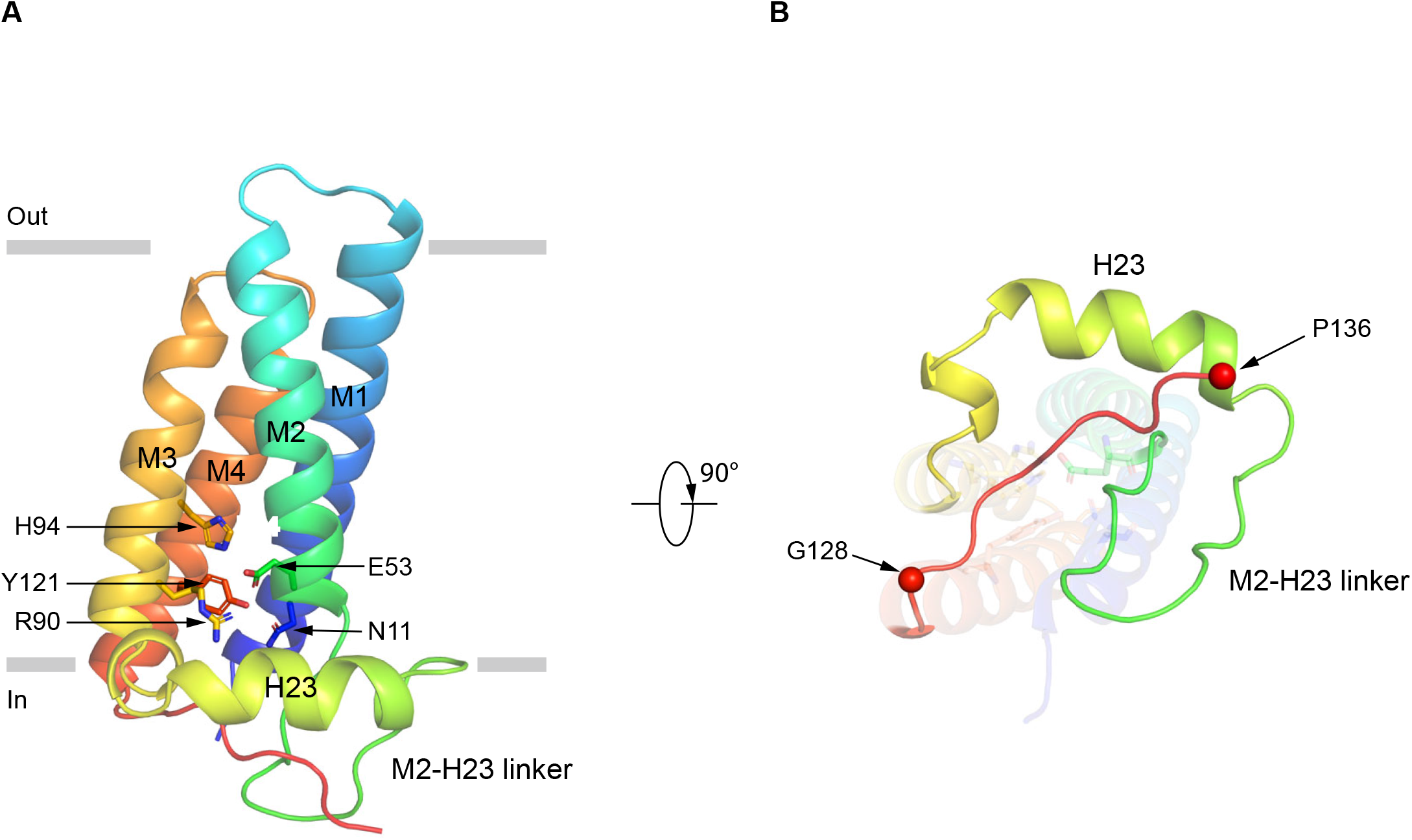
The structure of human p22 A, Structure of p22 in cartoon representation, and is colored in the rainbow pattern (N-terminus is blue and C-terminus is red). The phospholipid bilayer is shown as gray layers. B, The 90°-rotated bottom view of A. The 128-136 segment is colored in red. Cα positions of G128 and P136 are indicated as spheres.

### Interface between NOX2 and p22

The TMD of p22 makes extensive interactions with the TMD of NOX2, with a 9,618 Å^2^ interaction interface (Figure 4A-C). The assembly between NOX2 and p22 is mainly mediated by hydrophobic interactions, which involve M1 and M4 of p22 and M3, M4, and M5 of NOX2 (Figure 4D). Specifically, I4, W6, M8, W9, I20, V27, F33 on M1, L105, I108, L109, A112, I116 on M4 of p22 and I117, L120, F121 on M3, L167, A170, V180, L184, I187, L188 on M4, I197, Y201 on Loop D, V204 on M5 of NOX2 form two complementary shapes in the TMD of p22 and NOX2 for complex assembly (Figure 4D). These residues are conserved in NOX1-4 and p22 (Figure supplement 4-5). The tight packing between NOX2 and p22 supports their close association.

**Fig. 4.**
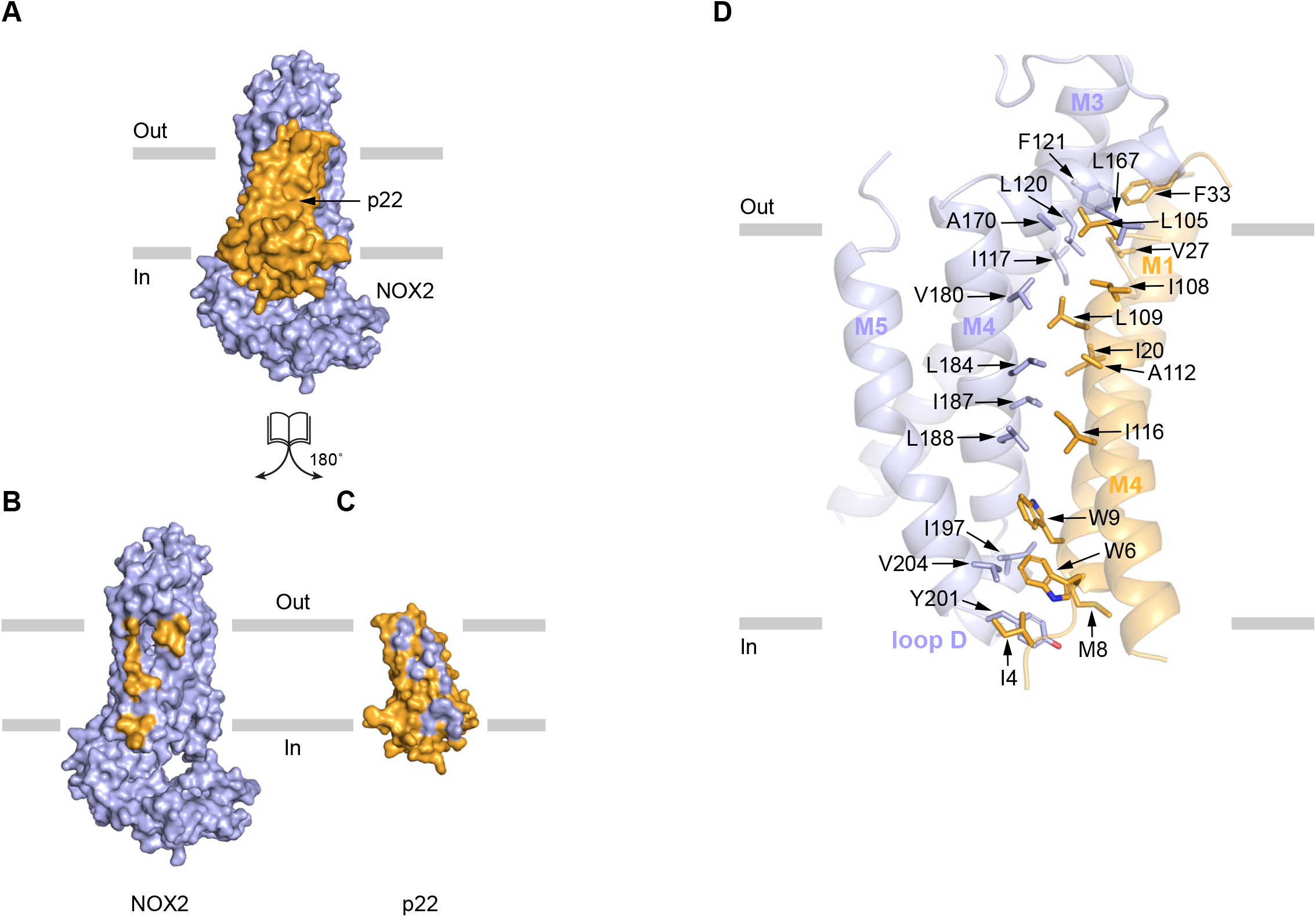
The interaction between NOX2 and p22 A-C, The open-book view of the interface between NOX2 and p22 in surface representation. NOX2 and p22 are colored in light blue and orange respectively in (A). Residues on NOX2 interacting with p22 are colored in orange in (B). Residues on p22 interacting with NOX2 are colored in light blue in (C). The phospholipid bilayer is shown as gray layers. D, The interface between NOX2 and p22 in cartoon representation. Residues participating in the interaction between NOX2 and p22 are shown as sticks.

### Conformational differences of DH between NOX2 and DUOX1

The poor local map quality of DH in the consensus refinement suggests its flexibility relative to TMD. Although after focused refinement, the map of FBD was improved, the map of NBD remained poor (Figure supplement 2E and F), indicating NBD is more flexible than FBD. Since the current structure represents the NOX2-p22 complex in an inactive resting state, we sought to understand how the complex is activated. Because the electron transfer pathway from DH to TMD (from FAD to inner haem) in the structure of DUOX1 in the high-calcium state ^7^ is in a reasonable range for efficient electron relay, we used it as the reference for structural comparison.

Alignment using their transmembrane domains shows that there is an obvious displacement of the DH domain of NOX2 (Figure 5A-B). The center of mass of FBD domains moves 9.5 Å (Figure 5C). There is a concomitant 13.8 Å movement of FAD which is bound on FBD (Figure 5D). Although the DH of NOX2 is still below and adjacent to its TMD, the orientation of FAD is not in a proper position for efficient electron transfer to inner haem of NOX2.

**Fig. 5.**
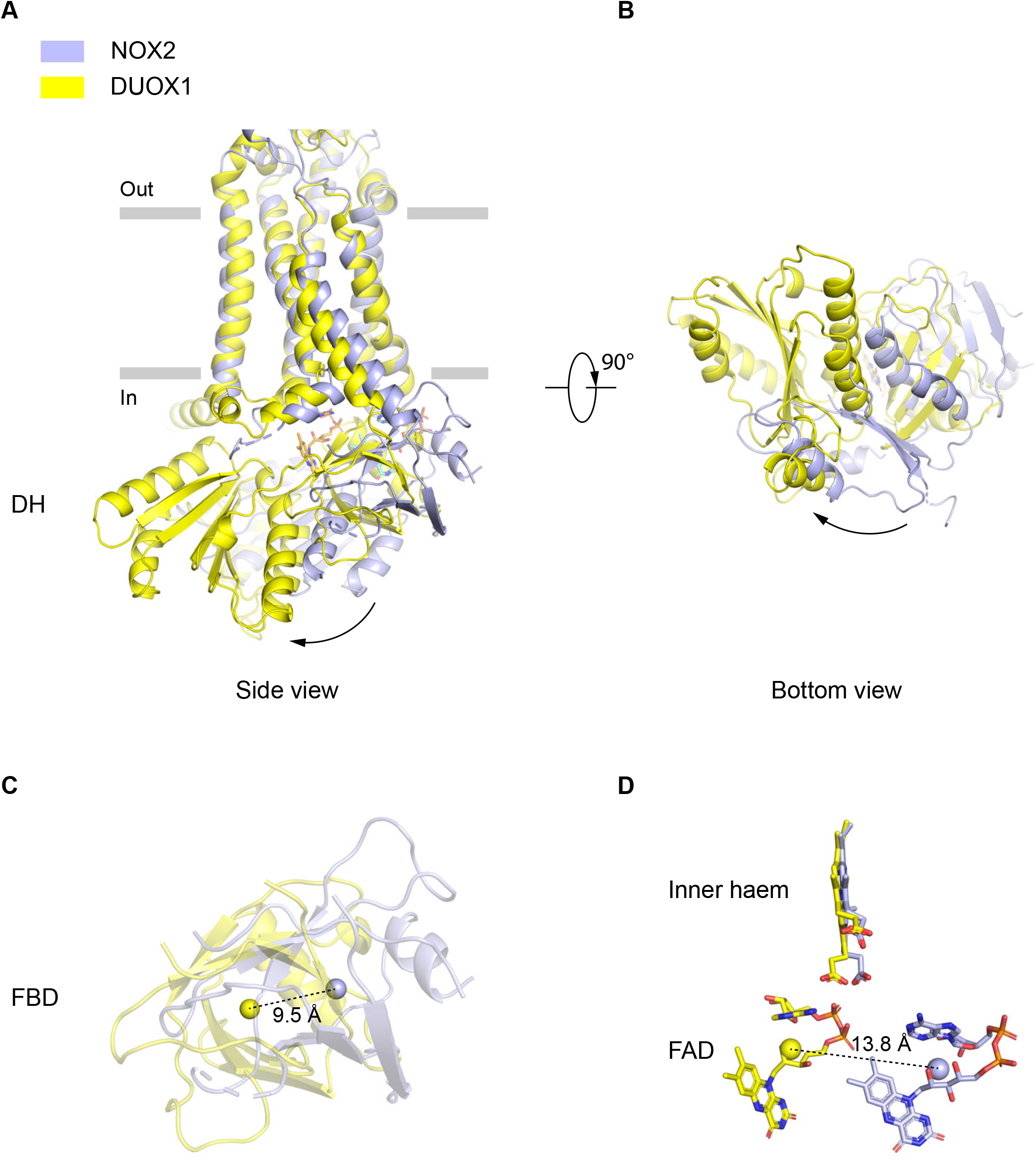
The movements of the DH domain of NOX2 compared with DUOX1 A, Side view of the TMD and dehydrogenase (DH) domain. NOX2 and DUOX1 are aligned using their transmembrane domain (TMD) in cartoon representation. NOX2 is colored in light blue and DUOX1 is colored in yellow. B, Bottom view of DH. The change of DH from NOX2 to DUOX1 was indicated by an arrow. C, Side view of FAD binding domain (FBD). The centers of mass of NOX2 and DUOX1 are shown as spheres respectively. The distance between centers of mass is shown in dashes. D, FAD and inner haem. The centers of mass of FAD in NOX2 and DUOX1 are shown as spheres respectively.

## Discussion

Although the common architecture of the NOX enzyme core and how they transfer electrons from intracellular NADPH to extracellular oxygen are emerging from previous structural studies ^5-7^, the structure of the essential auxiliary subunit p22 and how p22 assembles with NOX1-4 remain elusive prior to our current study. Here we provide the structure of the human NOX2-p22 heterodimer in the resting state at high resolution. Since the residues on NOX2 that interact with p22 are mostly conserved in NOX1-4 (Figure supplement 4), we speculate that NOX1, NOX3, and NOX4 likely interact with p22 in the same manner. The interaction mode between NOX2 and p22 is drastically distinct from that of DUOX1 and DUOXA1 (Figure supplement 7).DUOXA1 interacts with DUOX1 mainly through extracellular domains, including CLD of DUOXA1 and PHD of DUOX1. Moreover, DUOXA1 packs at the peripheral of M5 and M6 of DUOX1 (Figure supplement 7), whereas p22 packs against M3 and M4 of NOX2 (Figure supplement 7). Interestingly, the position of p22 is similar to the M0 of DUOX1 (Figure supplement 7). The C-terminus of M0 connects the intracellular regulatory PHLD and EF domains ^7^ and the C-terminus of p22 is the hub for the assembly of cytosolic factors for NOX2 activation. Therefore, both DUOX1 M0 and p22 recruit important regulatory modules of NOX, suggesting that this position might be a common binding site for the regulatory component of the NOX enzyme.

The NOX2-p22 structure presented here is in resting state. The edge-to-edge distances between haems and the electron-rely F215 are in a suitable range for electron transfer. But the distance between inner haem and FAD (5.6Å) is much larger than that of DUOX1 in the high calcium condition (3.9 Å), suggesting inefficient electron transfer from FAD to the transmembrane domain. Moreover, the NBD of DH is highly mobile, indicating weak interactions between NBD and other adjacent structural modules such as FBD and TMD of NOX2. Structural comparison between NOX2-p22 and DUOX1 in the high calcium condition shows the DH of DUOX1 docks at the bottom of TMD and has an obvious displacement compared to the DH of NOX2.

Therefore, we designate the current conformation of NOX2 DH as “undocked” and we speculate during the activation of NOX2, the cytosolic factors might likely stabilize the DH of NOX2 in the “docked” conformation which is similar to that observed in the activated DUOX1 in the high-calcium state to support the redox activity of NOX2 (Figure 6).

**Fig. 6.**
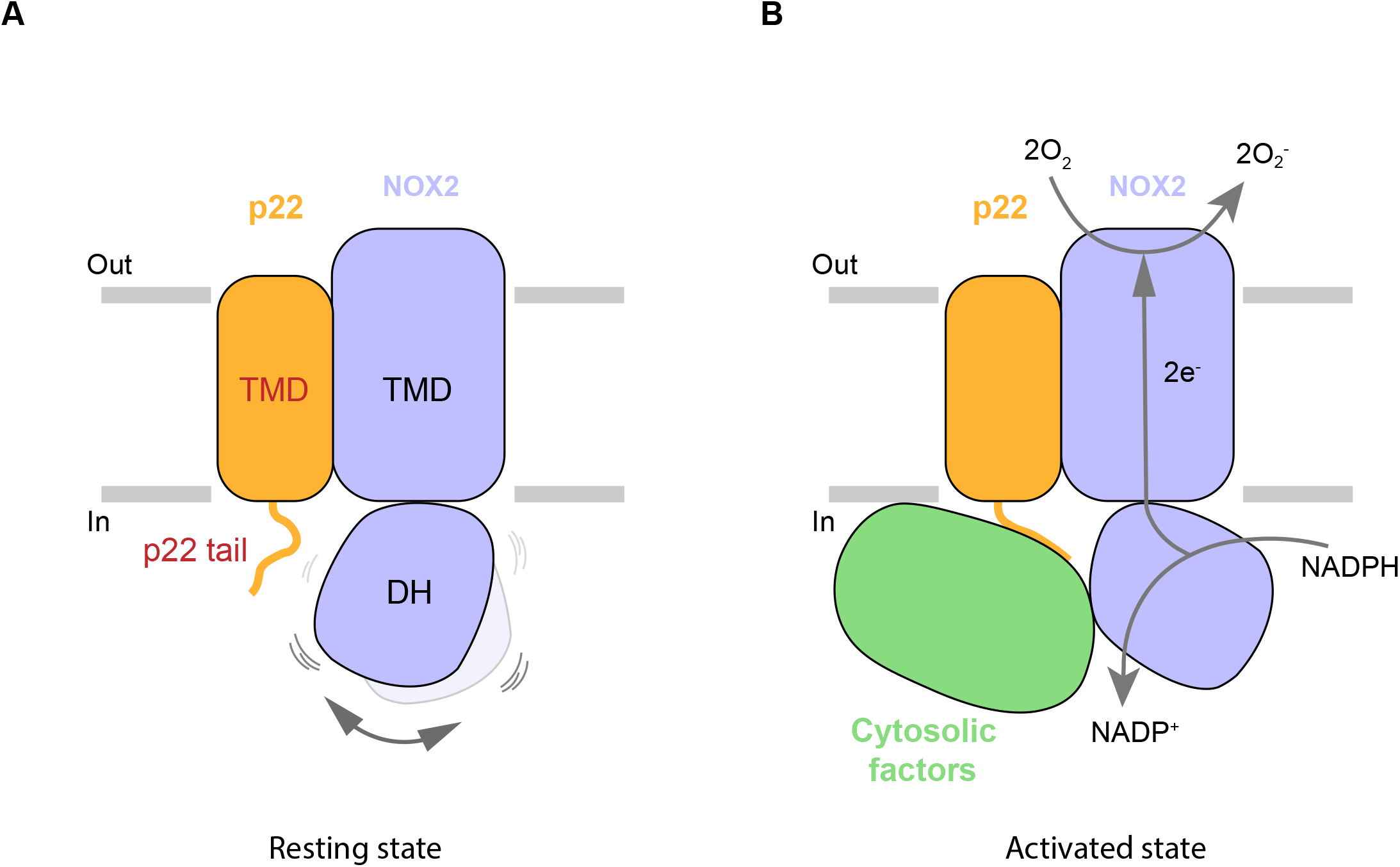
Hypothetic model for NOX2 activation A, Resting state of the phagocyte NADPH oxidase. NOX2, p22, and cytosolic factors are shown as cartoon and colored the same as Fig. 1A. B, Activated state of the phagocyte NADPH oxidase. The electron transfer pathway is indicted with gray arrows.

### Methods Cell culture

Sf9 insect cells (Thermo Fisher Scientific) were cultured in Sf-900 III serum-free medium (Gibco) or SIM SF serum-free medium (Sino Biological) at 27 °C. HEK293F suspension cells (Thermo Fisher Scientific) were cultured in FreeStyle 293 medium (Thermo Fisher Scientific) supplemented with 1% fetal bovine serum (FBS) at 37 °C with 6% CO_2_ and 70% humidity. The cell lines were routinely checked to be negative for mycoplasma contamination but have not been authenticated.

### Enzymatic assay

The superoxide anion-generating activity of NOX2-p22 complex was determined using the Amplex Red assay ^22^. The concentrations of H_2_O_2_ solution were determined by measuring UV-Vis absorbance at 240 nm with spectrophotometer (Pultton) and calculated using molar extinction coefficient of 43.6 M^−1^ cm^−1^. The concentration of H_2_O_2_ solution was further validated by reacting with Amplex Red to generate resorufin which has ε_571_=69,000 M^−1^ cm^−1 22^. Then the H_2_O_2_ solution with known concentration was used to calibrate the standard resorufin fluorescence curve (excitation, 530 nm; emission, 590 nm) measured using a Microplate Reader (Tecan Infinite M Plex) at 30 °C.

The superoxide anion-generating reaction of the membrane fraction containing NOX2-p22 complex was performed at 30 °C in 0.15 ml of HBS (20 mM HEPES, pH7.5, 150 mM NaCl) with 1 mM MgCl_2_, 1 mM EGTA, 10 μM FAD, 50 μM NADPH, 25 μM Amplex Red, 0.067 mg/ml horseradish peroxidase and 0.0576 mg/ml SOD. 33 nM Trimera and 10 μg/ml DPI were added as indicated. Progress of the reactions was monitored continuously by following the increase of the resorufin fluorescence, and the initial reaction rates were obtained by fitting the curve with linear equation. The data was processed with Microsoft Excel-2013 and GraphPad Prism 6.

### Protein expression, purification and nanodisc reconstitution

Gene encoding human p47^phox^ (1-286), human p67^phox^ (1-226), and full length human Rac1Q61L were fused together with the In-Fusion HD Cloning Kit (Takara Bio) and linked into a pET-based vector containing N-terminal 6×His-tag. The sequence of the linker between p47^phox^ and p67^phox^ was AAASTGGGSS, and the sequence of the linker between p67^phox^ and Rac1Q61L was GGGGGG. His6-Trimera protein was expressed in BL21 (DE3) and purified using Talon resin and HiTrap Q HP anion exchange chromatography column (GE Healthcare).

Mouse anti-human NOX2 monoclonal antibody 7D5 ^12^was purchased from MBL Life science. 7D5 Fab was released by papain digestion. Gene encoding TP1170 ^13^ was synthesized and cloned into a pET-based vector containing N-terminal GFP-tag and strep tag. GFP-strep-TP1170 protein was purified from Rosetta-gami2 (DE3).

Human p22 was fused with C-terminal GFP tag and strep tag and cloned into the pBMCL1 vector to generate pBMCL1-p22 ^23^. Human NOX2 was cloned into a modified BacMam vector ^24^ and linked into pBMCL1-p22 using LINK method^25^. The corresponding BacMam virus was made using sf9 cells ^7^. For protein expression, HEK293F cells cultured in Free Style 293 medium at a density of 2.5× 10^6^ cells per ml were infected with 10% volume of P2 virus. Sodium butyrate (10mM) and 5-Aminolevulinic acid hydrochloride (200μM) was added to the culture 12 h after transfection to promote protein expression, and the cells were transferred to a 30 °C incubator for another 36 h before harvesting. Cells were collected by centrifugation at 3,999×g (JLA-8.1000, Beckman Coulter) for 10 min at 4 °C, and washed with TBS buffer (20 mM Tris pH 8.0 at 4 °C, 150 mM NaCl) containing 2 μg/ml aprotinin, 2 μg/ml pepstatin, 2 μg/ml leupeptin. The cells were then flash-frozen and stored at -80 °C. For purification, cell pellet were resuspended in Buffer A [20 mM Hepes pH 7.5, 150 mM NaCl, 2 μg/ml aprotinin, 2 μg/ml pepstatin, 2 μg/ml leupeptin, 20% (v/v) glycerol, 1mM DTT] containing 1 mM phenylmethanesulfonyl fluoride (PMSF), 2 mM EGTA, 10 mM MgCl2, 10 μg/ml DNAse, 1% (w/v) DDM and 0.1% (w/v) CHS. The mixture was stirred for 1 hour at 4 °C to solubilize membrane proteins. The insoluble debris was removed by centrifugation at 193,400×g (Ti50.2, Beckman Coulter) for 40 min. Subsequently, the supernatant was loaded onto 4 ml Streptactin Beads 4FF (Smart Lifesciences) column and washed with 50 ml wash buffer 1 (Buffer A supplemented 0.5 mM DDM, 10 mM MgCl_2_ and 2 mM adenosine triphosphate (ATP)) to remove contamination of heat shock proteins. Then, the column was extensively washed with 50 ml wash buffer 2 (Buffer A supplemented with 0.5 mM DDM). The target protein was eluted with Buffer A supplemented 0.5 mM DDM, and 8.5 mM D-desthiobiotin (IBA). The eluate was loaded onto HiTrap Q HP (GE Healthcare) and the NOX2-p22 complex was separated from aggregates with a linear gradient from 0 mM NaCl to 1000 mM NaCl in buffer containing 20 mM Hepes pH 7.5 at 4 °C, 0.5 mM DDM. The fractions containing the NOX2-p22 complex were collected for nanodisc reconstitution. NOX2-p22 complex was mixed with MSP2X and L-α-phosphatidylcholine from soybean (PC, solubilized in 2% (w/v) OG) at a molar ratio of protein: MSP2X: PC = 1: 4: 800. The mixture was allowed to equilibrate for 1 h at 4 °C, and then Bio-beads SM2 (Bio-Rad) was added to initiate the reconstitution with constant rotation at 4 °C. Bio-beads SM2 were added to the mixture three times within 15 h to gradually remove detergents from the system. Afterwards, the proteins that were not reconstituted in lipid nanodisc was removed by centrifugation at 86,600×g for 30 min in TLA 100.3 rotor (Beckman Coulter). The supernatant containing NOX2-p22 complex reconstituted in lipid nanodisc was loaded onto the 1 ml Streptactin Beads 4FF column to remove empty nanodisc. The elution from Streptactin Beads 4FF column was concentrated and subjected onto a Superose 6 increase 10/300 GL column (GE Healthcare) in buffer that contained 20 mM HEPES pH 7.5, 150 mM NaCl. The peak fractions corresponding to the NOX2-p22 complex in lipid nanodisc were collected. The GFP tag at the C-terminus of p22 was cleaved off by Prescission Protease digestion. Then 7D5 Fab and GFP-strep-TP1170 protein was added into NOX2-p22 nanodisc sample and the mixture was subjected to Streptactin Beads 4FF to remove excess p22 and free Fc fragment. NOX2-p22-7D5-TP1170 quaternary complex was eluted with D-desthiobiotin, concentrated and subjected to Superose 6 increase 10/300 GL column. Peak fractions containing the complex were pooled for cryo-EM sample preparation. Concentration of NOX2 was estimated with absorbance at 415 nm (ε_415_=0.262 μM^−1^ cm^−1^)^26^.

### Cryo-EM sample preparation and data collection

To obtain a homogenous orientation distribution of particles, the surface of Quantifoil Au 300 mesh R 0.6/1.0 grids were coated with graphene oxide (GO grids) ^14^ or graphene (G grids)^15^.Aliquots of 2.5 µl of NOX2-p22 protein in nanodisc at a concentration of approximately 0.12 mg/ml in the presence of 0.5 mM FAD, 1 mM NADPH and 0.5 mM fluorinated octyl-maltoside (FOM, Anatrace) were applied to the grids. After 60 s incubation on the grids at 4 °C under 100% humidity, the grids were then blotted for 4 s using a blot force of 4, and then plunge-frozen into liquid ethane using a Vitrobot Mark IV (Thermo Fisher Scientific). The grids were transferred to a Titan Krios electron microscope (Thermo Fisher) operating at 300 kV and a K2 Summit direct detector (Gatan) mounted post a quantum energy filter (slit width 20 eV). SerialEM-3.6.11 was used for automated data collection. Movies from dataset of NOX2-p22 were recorded in super-resolution mode and a defocus range of -1.5 µm to -2.0 μm with a nominal magnification of 165,000×, resulting in a calibrated pixel size of 0.821 Å, Each stack of 32 frames was exposed for 8 s, with an exposing time of 0.25 s per frame at a dose rate of 4.7 electrons per Å^−2^ per second.

### Image processing

A total of 1,890 and 2,059 micrographs of the NOX2-p22 complex on graphene oxide grid and graphene grid were collected, separately. The beam-induced drift was corrected using MotionCor2 ^27^ and binned to a pixel size of 1.642 Å. Dose-weighted micrographs were used for CTF estimation by Gctf^28^. A total of 206,504 and 355,928 particles were auto-picked using Gautomatch-0.56 (developed by K. Zhang) or Topaz-0.2.3 ^29^. Two rounds of 2D classification in Relion ^30^ were performed to remove ice contaminants and aggregates, yielding 14,287 and 27,177 particles and used as seeds for seed-facilitated classification ^31^. After that two datasets were combined and duplicated particles were removed, and 208,906 particles were retained and subjected to multi-reference 3D classification using resolution gradient was performed ^31^. The resulting particles of selected classes were subjected to non-uniform refinement and local refinement to generate a consensus map at 3.3 Å resolution ^32^. To further improve the local map quality of extracellular TP1170-constant region of Fab fragment and intracellular DH, these regions were subjected to focused refinement in cryoSPARC to generate two local maps. These two local maps were merged into a full consensus map to generate a composite map for model building. All resolutions were estimated using the gold-standard Fourier shell correlation 0.143 criterion. Local resolution was calculated using cryoSAPRC^30^.

### Model building

Predicted models of NOX2 and p22 were downloaded from Alpha Fold 2 database ^33^. Fab was modeled as poly-alanine. Model of TP1170 was modeled using SWISS-MODEL^34^. Individual models were docked into the map using Chimera ^35^. The model were iteratively manually rebuilt using Coot^36^ according to the map and refined using Phenix ^37,38^.

### Quantification and statistical analysis

The number of independent reactions (N) and the relevant statistical parameters for each experiment (such as mean or standard deviation) are described in the figure legends. No statistical methods were used to pre-determine sample sizes.

## Data availability

Cryo-EM maps and atomic coordinate of the NOX2-p22-7D5-TP1170 complex have been deposited in the EMDB and PDB under the ID codes EMDB: EMD-34389 and PDB: 8GZ3.

## Acknowledgements

We greatly thank Miao Wei for providing the GFP-strep-TP1170 protein. We thank Xiaoyu Liu and Yiting Shi for illustration. Cryo-EM data collection was supported by Electron microscopy laboratory and Cryo-EM platform of Peking University with the assistance of Xuemei Li, Zhenxi Guo, Changdong Qin, Xia Pei, and Guopeng Wang. Part of the structural computation was also performed on the Computing Platform of the Center for Life Science and High-performance Computing Platform of Peking University. We thank the National Center for Protein Sciences at Peking University in Beijing, China for assistance with negative stain EM. The work is supported by grants from the National Natural Science Foundation of China (91957201, 31870833, and 31821091 to L.C.; 52021006 to H.P.), Beijing National Laboratory for Molecular Sciences (BNLMS-CXTD-202001 to H.P.) the National Key Research and Development Program of China, and Center For Life Sciences (CLS to L.C.).

## Author contributions

L.C. initiated the project and wrote the manuscript draft. R.L., K.S., J-X.W., and X. Geng screened constructs and established the methods for activity assay. R.L. and K.S. purified proteins. L.Z., X.Gao, and H.P. prepared the graphene coated cryo-EM grids. R.L. prepared the cryo-EM samples, collected the cryo-EM data, processed data, built and refined the model. All authors contributed to the manuscript preparation.

## Figure Legends

**Fig. S1.**
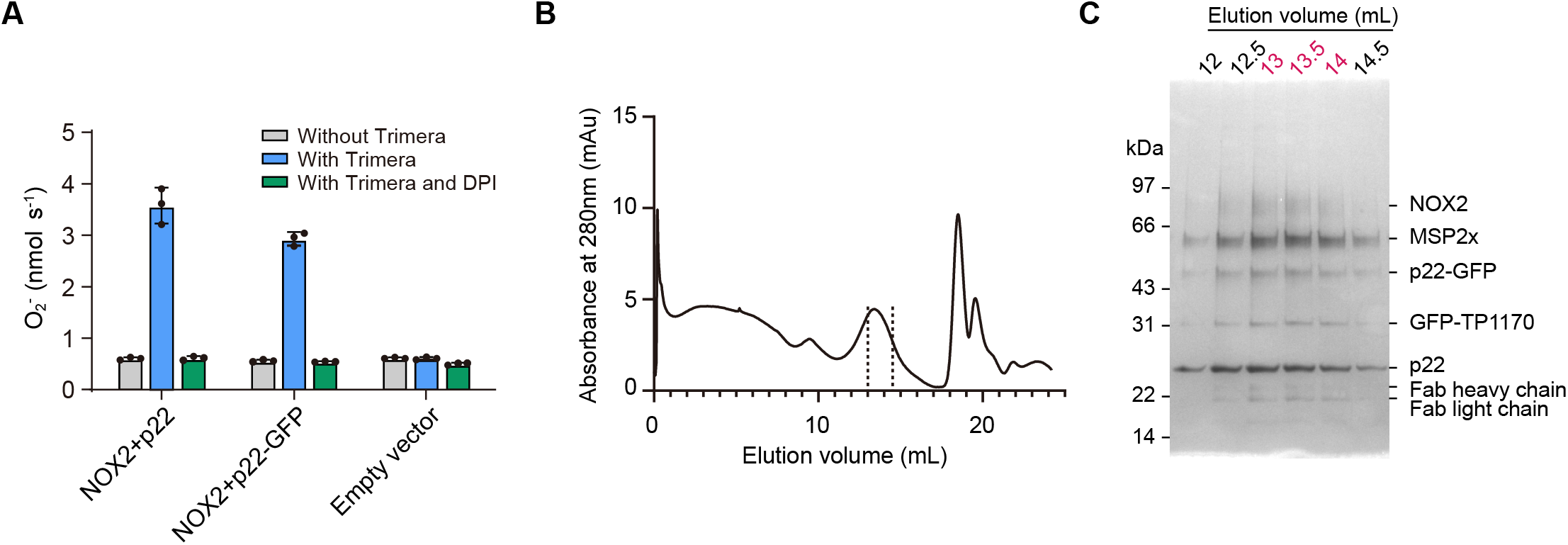
Protein purification A, The activity of the NOX2-p22 complex in crude cell membrane measured using Amplex Red assay. Data are shown as means ± standard deviations; N = 3 technical replicates. B, Size-exclusion chromatography profile of the NOX2-p22-7D5-TP1170 complex in nanodisc. Fractions between dashes were used for cryo-EM sample preparation. C, SDS-PAGE gel of purified NOX2-p22-7D5-TP1170 protein in the nanodisc. Fractions with numbers in red were used for cryo-EM sample preparation. For uncropped gel, please see Figure 1-Figure supplement 1-Source Data 1.

**Fig. S2.**
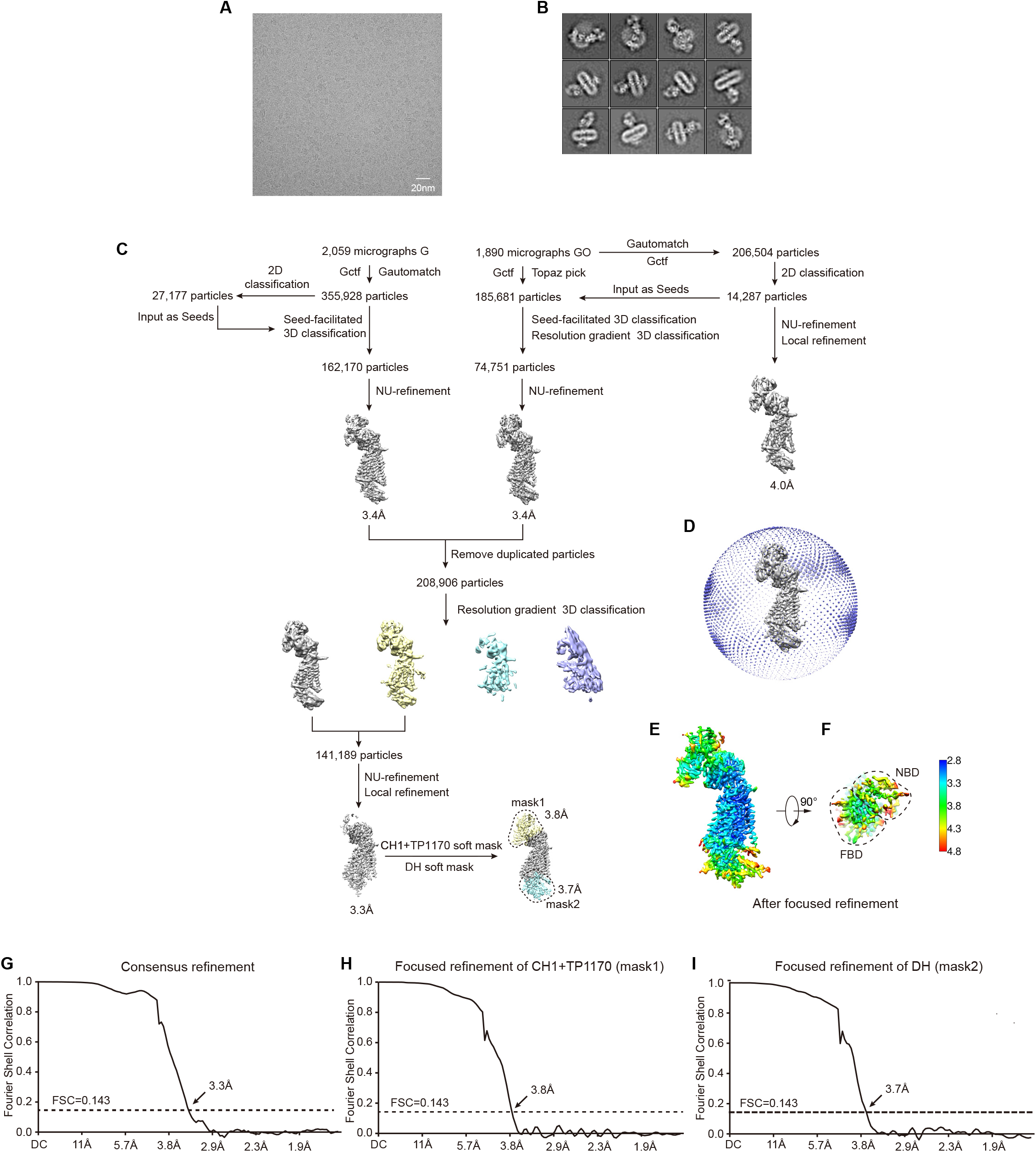
Image processing A, Representative raw micrograph of NOX2-p22-7D5-TP1170 complex in nanodisc. B, 2D class averages of NOX2-p22-7D5-TP1170 complex. C, Workflow of the cryo-EM data processing. D, Angular distributions of the final consensus reconstruction. E, Local resolution of the NOX2-p22-7D5-TP1170 complex after focused refinement. F, Gold-standard Fourier shell correlation curves of the final consensus refinement. G, Gold-standard Fourier shell correlation curves of the focused refinement of the TP1170-constant region of Fab fragment (CH1+TP1170) using mask 1. H, Gold-standard Fourier shell correlation curves of the focused refinement of the DH domain using mask 2.

**Fig. S3.**
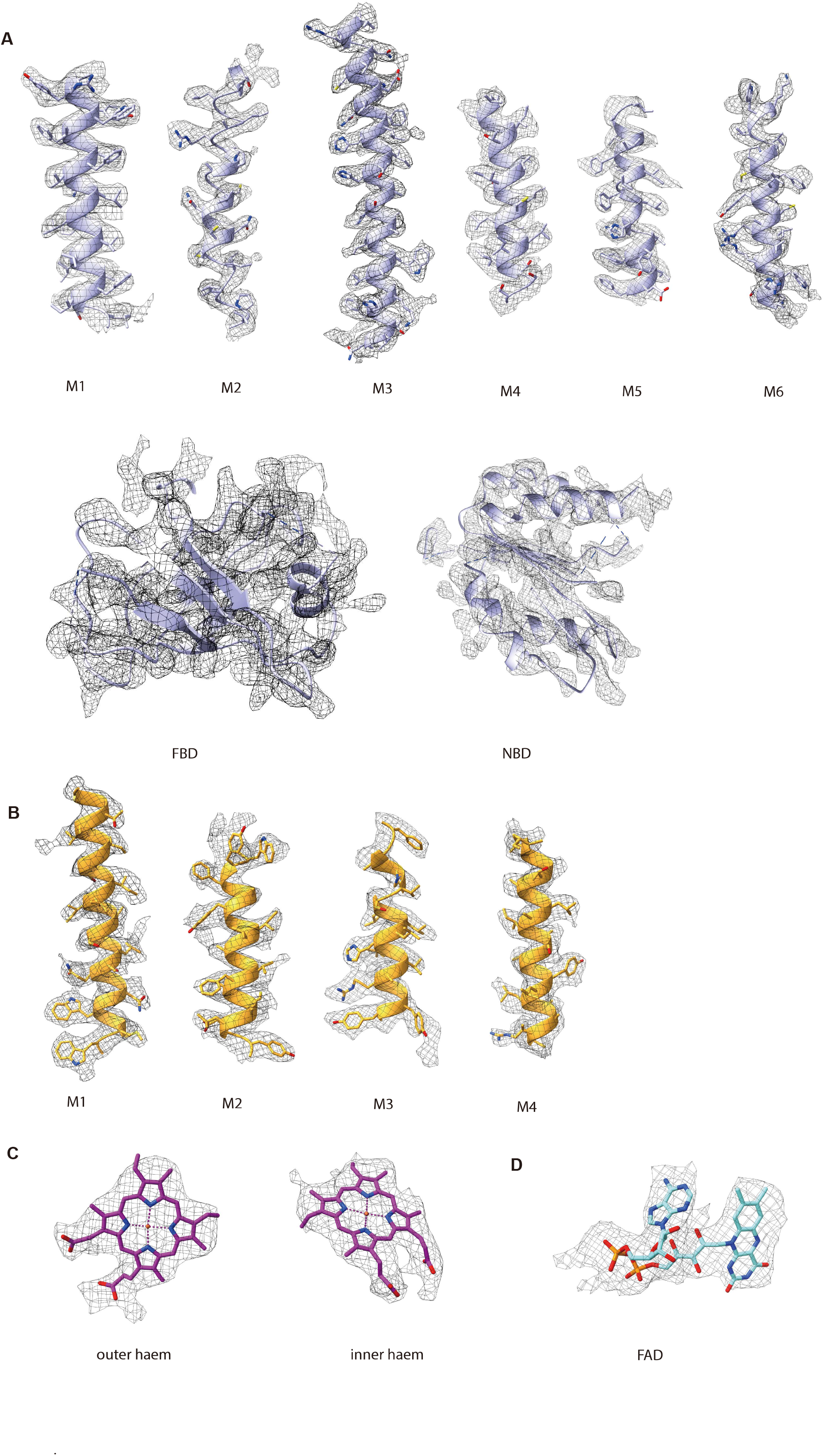
Representative local electron density maps A, Local electron density maps of NOX2. B, Local electron density maps of p22. C. Local electron density maps of haems bound in NOX2. D. Local electron density map of FAD bound in FBD of NOX2.

**Fig. S4.**
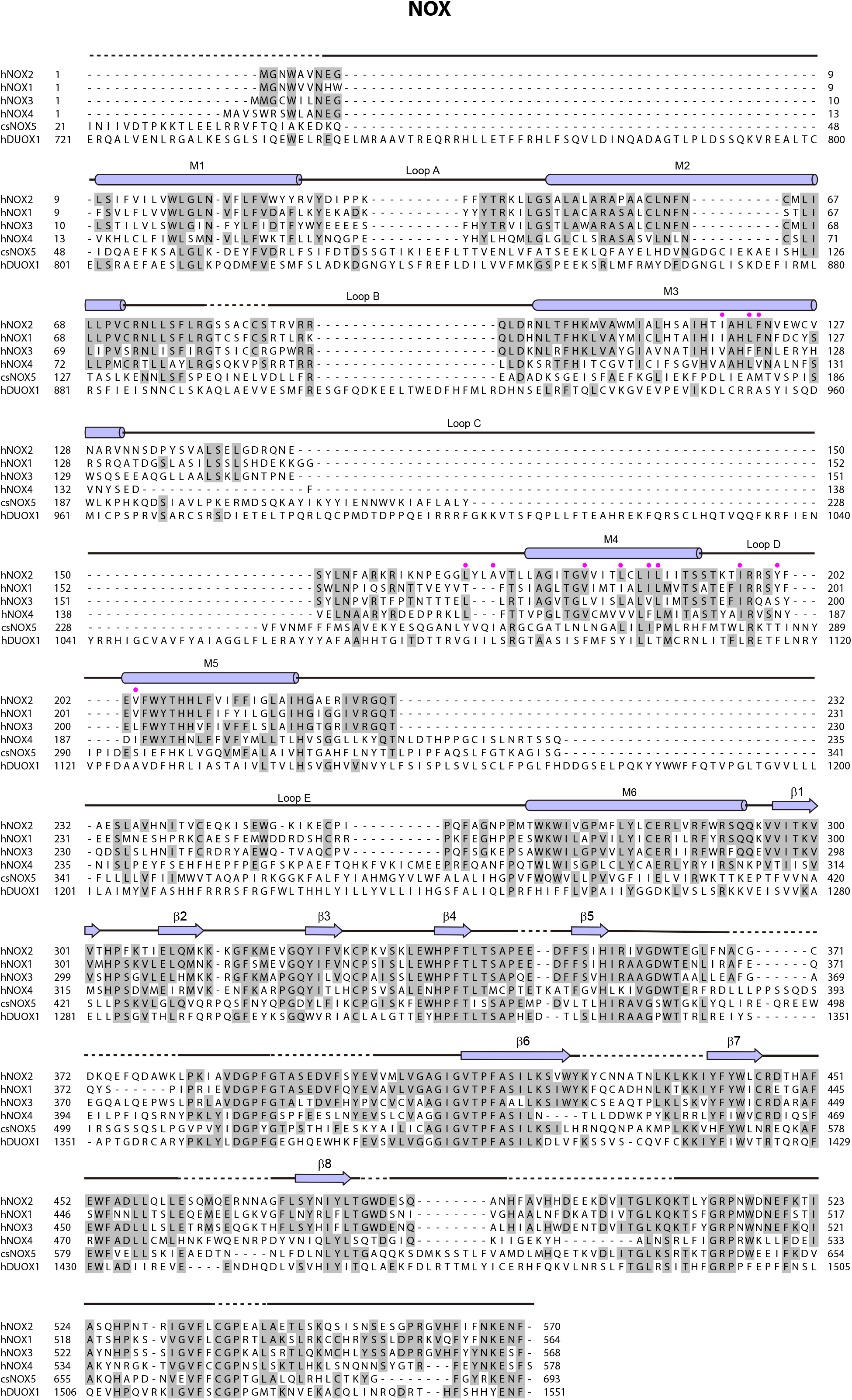
Sequence alignment of the NOX subunit The sequences of the *Homo sapiens* NOX2 (hNOX2, UniProtKB: P04839), *Homo sapiens* NOX1 (hNOX1, UniProtKB: Q9Y5S8), *Homo sapiens* NOX3 (hNOX3, UniProtKB: Q9HBY0), *Homo sapiens* NOX4 (hNOX4, UniProtKB: Q9NPH5), *Cylindrospermum stagnale* NOX5 (csNOX5, UniProtKB: K9WT99) and *Homo sapiens* DUOX1 (hDUOX1, UniProtKB: Q9NRD9) were aligned. The sequence alignment of Fig. S4 and S5 are all shown as follows: Conserved residues are highlighted in gray; Secondary structures are indicated as cylinders (α helices), arrows (β sheets), and lines (loops); Unmodeled residues are indicated as dashed lines; The color of arrows and cylinders are the same as in Fig. 1I. Residues interact with p22 are indicated as pink-filled circles.

**Fig. S5.**
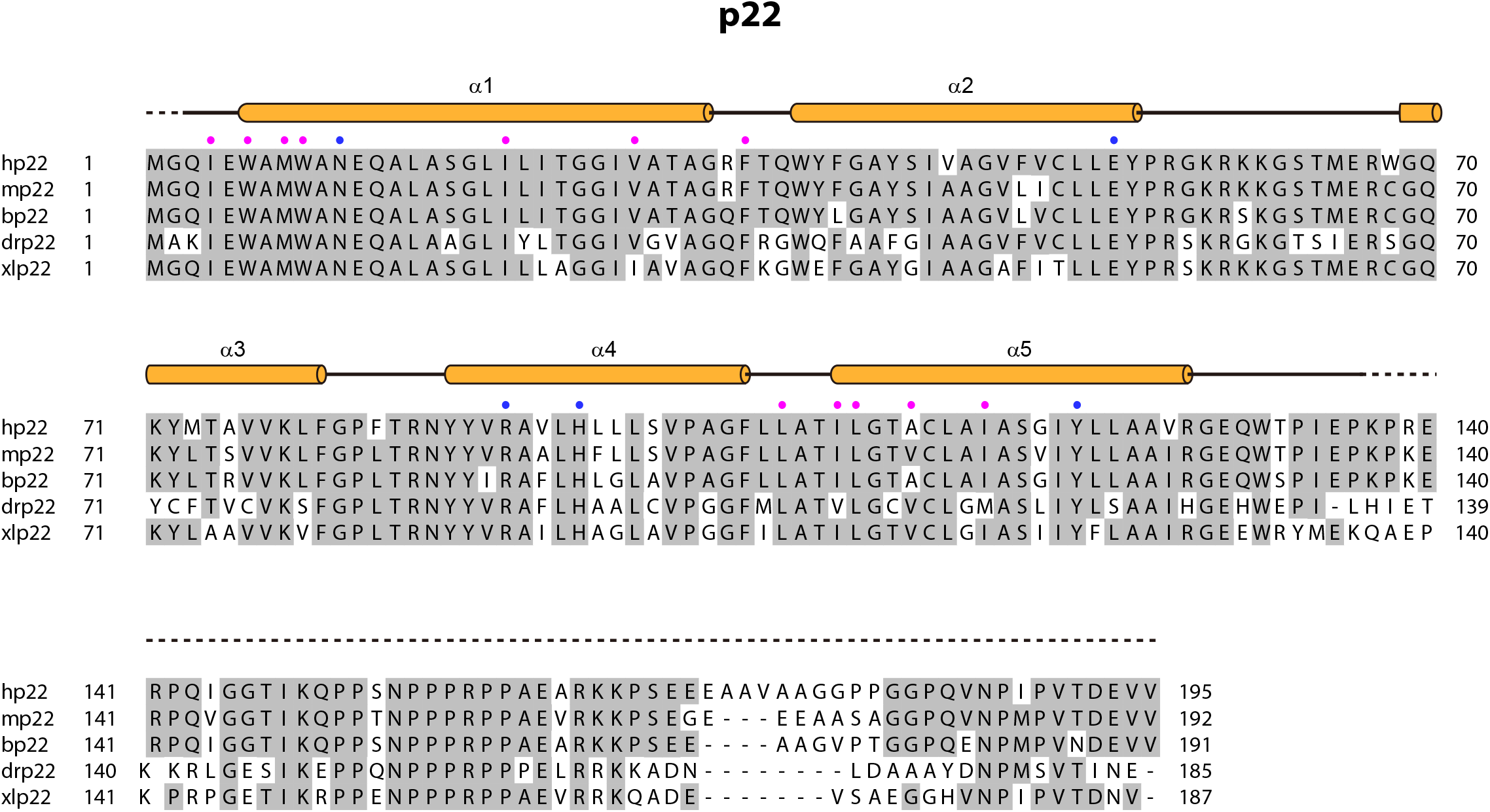
Sequence alignment of p22 subunit The sequences of the *Homo sapiens* p22 (hp22, UniProtKB: P13498), *Mus musculus* p22 (mp22, UniProtKB: Q61462), *Bos taurus* p22 (bp22, UniProtKB: O46521), *Danio rerio* p22 (drp22, UniProtKB: Q6PH62) and *Xenopus laevis* p22 (xlp22, UniProtKB: Q6AZJ1) were aligned. Residues interacting with NOX2 are indicated as pink-filled circles. The blue-filled circles indicate the residues that form a polar interaction network.

**Fig. S6.**
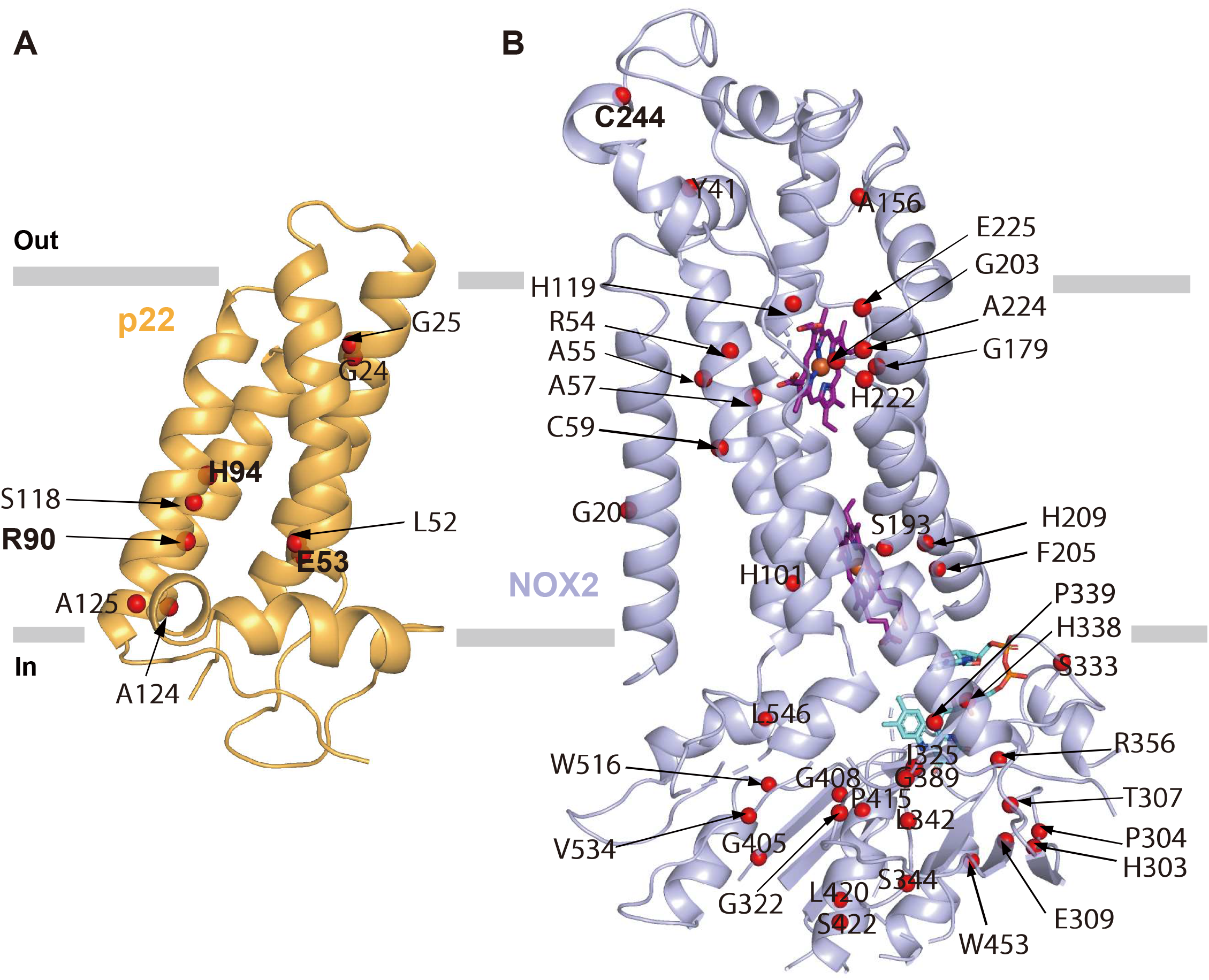
Locations of CGD mutations found in human patients A, Mutations of NOX2 found in CGD patients (annotated in UNIPROT) are mapped onto the structural model of NOX2. Cα positions of mutations are shown as red spheres, NOX2 is shown in a light blue cartoon. B, Mutations of p22 found in CGD patients (annotated in UNIPROT) are mapped onto the structual model of p22. Cα positions of mutations are shown as red spheres. p22 is shown in a bright orange cartoon. Mutations mentioned in the text are highlighted in bold.

**Fig. S7.**
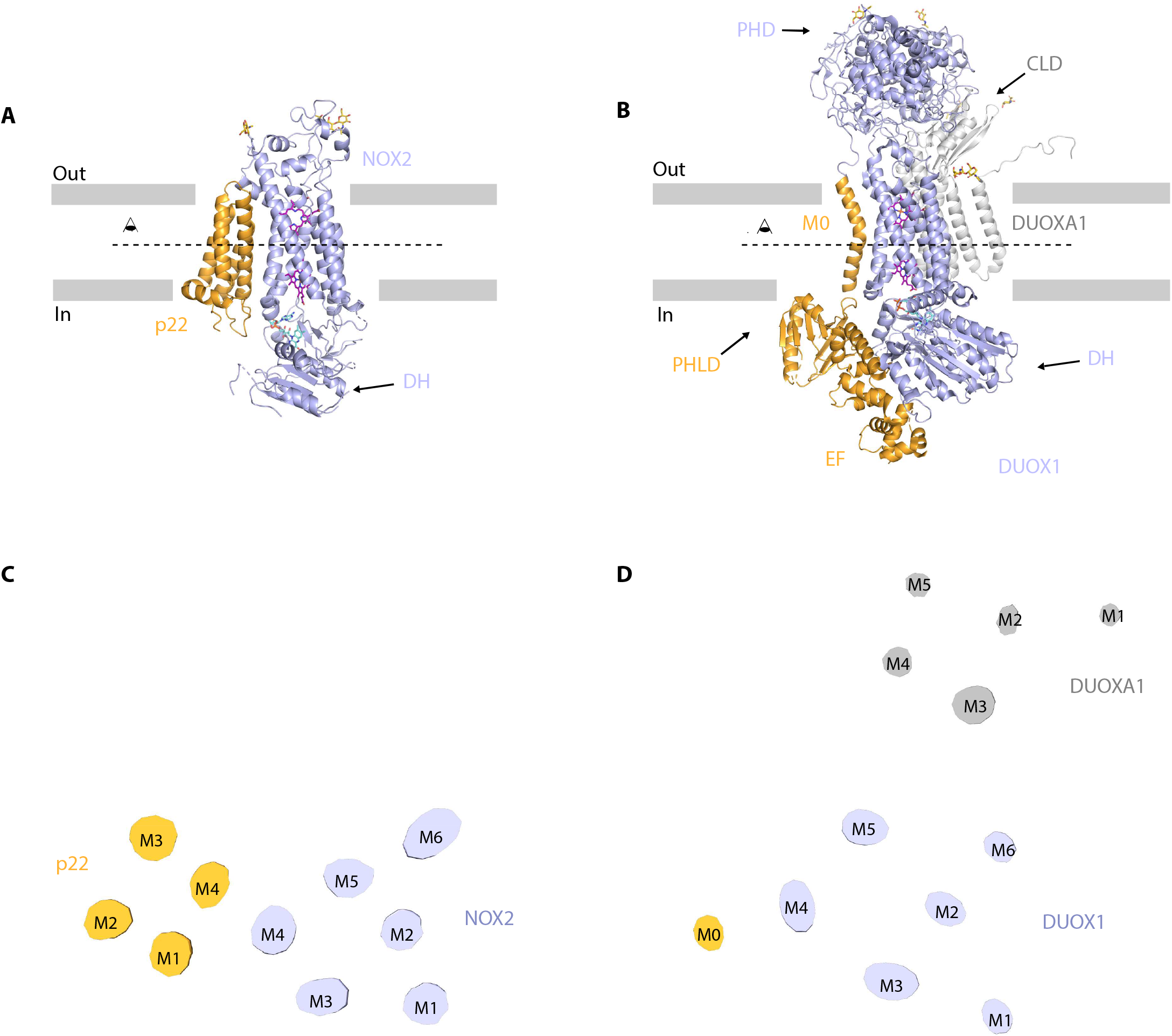
Distinct interaction mode between NOX2-p22 complex and DUOX1-DUOXA1 complex A, Structure of human NOX2 and p22 complex (PDB ID: 8GZ3) in cartoon representation. The colors of each individual domain are the same as in Fig.1I. The approximate boundaries of the phospholipid bilayer are indicated as gray thick lines. Sugar moieties, haems, and FAD are shown as gold, purple and aquamarine sticks, respectively. B, Structure of human DUOX1-DUOXA1 complex (PDB ID: 7D3F) in cartoon representation. PHD: peroxidase homology domain of DUOX1; PHLD: pleckstrin homology-like domain of DUOX1; EF: EF-hand calcium-binding module of DUOX1; DH: dehydrogenase domain of DUOX1; CLD: claudin-like domain of DUOXA1. The M0 helix, PHLD, and EF are colored in orange, and the other part of DUOX1 is colored in blue. The DUOXA1 is colored in grey. C, Top view of the cross-section of NOX2-p22 cryo-EM map on the transmembrane layer at the position indicated as a dashed line in A. For clarity, the cryo-EM map was low-pass filtered to 7 Å. D, Top view of the cross-section of DUOX1-DUOXA1 cryo-EM map on the transmembrane layer at the position indicated as a dashed line in B.

## References

1 Lambeth, J. D. & Neish, A. S. Nox enzymes and new thinking on reactive oxygen: a double-edged sword revisited. Annu. Rev. Pathol. 9, 119–145, doi:10.1146/annurev-pathol-012513-104651 (2014).

2 Bedard, K. & Krause, K. H. The NOX family of ROS-generating NADPH oxidases: physiology and pathophysiology. Physiol. Rev. 87, 245–313, doi:10.1152/physrev.00044.2005 (2007).

3 Winterbourn, C. C., Kettle, A. J. & Hampton, M. B. Reactive Oxygen Species and Neutrophil Function. Annu. Rev. Biochem. 85, 765–792, doi:10.1146/annurev-biochem-060815-014442 (2016).

4 Heyworth, P. G., Cross, A. R. & Curnutte, J. T. Chronic granulomatous disease. Curr. Opin. Immunol. 15, 578–584, doi:10.1016/s0952-7915(03)00109-2 (2003).

5 Magnani, F. et al. Crystal structures and atomic model of NADPH oxidase. Proc. Natl. Acad. Sci. U. S. A. 114, 6764–6769, doi:10.1073/pnas.1702293114 (2017).

6 Sun, J. Structures of mouse DUOX1-DUOXA1 provide mechanistic insights into enzyme activation and regulation. Nat. Struct. Mol. Biol., doi:10.1038/s41594-020-0501-x (2020).

7 Wu, J. X., Liu, R., Song, K. & Chen, L. Structures of human dual oxidase 1 complex in low-calcium and high-calcium states. Nat Commun 12, 155, doi:10.1038/s41467-020-20466-9 (2021).

8 Stasia, M. J. CYBA encoding p22(phox), the cytochrome b558 alpha polypeptide: gene structure, expression, role and physiopathology. Gene 586, 27–35, doi:10.1016/j.gene.2016.03.050 (2016).

9 Meijles, D. N., Howlin, B. J. & Li, J. M. Consensus in silico computational modelling of the p22phox subunit of the NADPH oxidase. Comput. Biol. Chem. 39, 6–13, doi:10.1016/j.compbiolchem.2012.05.001 (2012).

10 Forteza, R., Salathe, M., Miot, F. & Conner, G. E. Regulated hydrogen peroxide production by Duox in human airway epithelial cells. Am. J. Respir. Cell Mol. Biol. 32, 462–469, doi:10.1165/rcmb.2004-0302OC (2005).

11 Berdichevsky, Y., Mizrahi, A., Ugolev, Y., Molshanski-Mor, S. & Pick, E. Tripartite chimeras comprising functional domains derived from the cytosolic NADPH oxidase components p47phox, p67phox, and Rac1 elicit activator-independent superoxide production by phagocyte membranes: an essential role for anionic membrane phospholipids. J. Biol. Chem. 282, 22122–22139, doi:10.1074/jbc.M701497200 (2007).

12 Yamauchi, A. et al. Location of the epitope for 7D5, a monoclonal antibody raised against human flavocytochrome b558, to the extracellular peptide portion of primate gp91phox. Microbiol. Immunol. 45, 249–257, doi:10.1111/j.1348-0421.2001.tb02614.x (2001).

13 Pleiner, T., Bates, M. & Gorlich, D. A toolbox of anti-mouse and anti-rabbit IgG secondary nanobodies. J. Cell Biol. 217, 1143–1154, doi:10.1083/jcb.201709115 (2018).

14 Patel, A., Toso, D., Litvak, A. & Nogales, E. Efficient graphene oxide coating improves cryo-EM sample preparation and data collection from tilted grids. bioRxiv (2021).

15 Zhang, J. et al. Clean Transfer of Large Graphene Single Crystals for High-Intactness Suspended Membranes and Liquid Cells. Adv Mater 29, doi:10.1002/adma.201700639 (2017).

16 Rae, J. et al. X-Linked chronic granulomatous disease: mutations in the CYBB gene encoding the gp91-phox component of respiratory-burst oxidase. Am. J. Hum. Genet. 62, 1320–1331, doi:10.1086/301874 (1998).

17 Bolscher, B. G., de Boer, M., de Klein, A., Weening, R. S. & Roos, D. Point mutations in the beta-subunit of cytochrome b558 leading to X-linked chronic granulomatous disease. Blood 77, 2482–2487 (1991).

18 Patino, P. J. et al. Molecular analysis of chronic granulomatous disease caused by defects in gp91-phox. Hum. Mutat. 13, 29–37, doi:10.1002/(SICI)1098-1004(1999)13:1<29::AID-HUMU3>3.0.CO;2-X (1999).

19 de Boer, M. et al. Cytochrome b558-negative, autosomal recessive chronic granulomatous disease: two new mutations in the cytochrome b558 light chain of the NADPH oxidase (p22-phox). Am. J. Hum. Genet. 51, 1127–1135 (1992).

20 Rae, J. et al. Molecular analysis of 9 new families with chronic granulomatous disease caused by mutations in CYBA, the gene encoding p22(phox). Blood 96, 1106–1112 (2000).

21 Hossle, J. P., de Boer, M., Seger, R. A. & Roos, D. Identification of allele-specific p22-phox mutations in a compound heterozygous patient with chronic granulomatous disease by mismatch PCR and restriction enzyme analysis. Hum. Genet. 93, 437–442, doi:10.1007/BF00201671 (1994).

22 Zhou, M., Diwu, Z., Panchuk-Voloshina, N. & Haugland, R. P. A stable nonfluorescent derivative of resorufin for the fluorometric determination of trace hydrogen peroxide: applications in detecting the activity of phagocyte NADPH oxidase and other oxidases. Anal. Biochem. 253, 162–168, doi:10.1006/abio.1997.2391 (1997).

23 Guo, W., Wang, M. & Chen, L. A co-expression vector for baculovirus-mediated protein expression in mammalian cells. Biochem. Biophys. Res. Commun. 594, 69–73, doi:10.1016/j.bbrc.2022.01.056 (2022).

24 Li, N. et al. Structure of a Pancreatic ATP-Sensitive Potassium Channel. Cell 168, 101–110 e110, doi:10.1016/j.cell.2016.12.028 (2017).

25 Scheich, C., Kummel, D., Soumailakakis, D., Heinemann, U. & Bussow, K. Vectors for co-expression of an unrestricted number of proteins. Nucleic Acids Res. 35, e43, doi:10.1093/nar/gkm067 (2007).

26 Jesaitis, A. J., Riesselman, M., Taylor, R. M. & Brumfield, S. Enhanced Immunoaffinity Purification of Human Neutrophil Flavocytochrome B for Structure Determination by Electron Microscopy. Methods Mol. Biol. 1982, 39–59, doi:10.1007/978-1-4939-9424-3_3 (2019).

27 Zheng, S. Q. et al. MotionCor2: anisotropic correction of beam-induced motion for improved cryo-electron microscopy. Nat. Methods 14, 331–332, doi:10.1038/nmeth.4193 (2017).

28 Zhang, K. Gctf: Real-time CTF determination and correction. J. Struct. Biol. 193, 1–12, doi:10.1016/j.jsb.2015.11.003 (2016).

29 Bepler, T. et al. Positive-unlabeled convolutional neural networks for particle picking in cryo-electron micrographs. Nat. Methods 16, 1153–1160, doi:10.1038/s41592-019-0575-8 (2019).

30 Zivanov, J. et al. New tools for automated high-resolution cryo-EM structure determination in RELION-3. Elife 7, doi:10.7554/eLife.42166 (2018).

31 Wang, N. et al. Structural basis of human monocarboxylate transporter 1 inhibition by anti-cancer drug candidates. Cell 184, 370–383 e313, doi:10.1016/j.cell.2020.11.043 (2021).

32 Punjani, A., Zhang, H. & Fleet, D. J. Non-uniform refinement: adaptive regularization improves single-particle cryo-EM reconstruction. Nat. Methods 17, 1214–1221, doi:10.1038/s41592-020-00990-8 (2020).

33 Jumper, J. et al. Highly accurate protein structure prediction with AlphaFold. Nature 596, 583–589, doi:10.1038/s41586-021-03819-2 (2021).

34 Biasini, M. et al. SWISS-MODEL: modelling protein tertiary and quaternary structure using evolutionary information. Nucleic Acids Res. 42, W252–258, doi:10.1093/nar/gku340 (2014).

35 Pettersen, E. F. et al. UCSF Chimera--a visualization system for exploratory research and analysis. J Comput Chem 25, 1605–1612, doi:10.1002/jcc.20084 (2004).

36 Emsley, P., Lohkamp, B., Scott, W. G. & Cowtan, K. Features and development of Coot. Acta Crystallogr. D Biol. Crystallogr. 66, 486–501, doi:10.1107/S0907444910007493 (2010).

37 Adams, P. D. et al. PHENIX: a comprehensive Python-based system for macromolecular structure solution. Acta Crystallogr. D Biol. Crystallogr. 66, 213–221, doi:10.1107/S0907444909052925 (2010).

38 Afonine, P. V. et al. Real-space refinement in PHENIX for cryo-EM and crystallography. Acta Crystallogr D Struct Biol 74, 531–544, doi:10.1107/S2059798318006551 (2018).

